# Mechanical history shapes phenotypic plasticity and fibroblast-autonomous 3D remodeling in iPSC-derived cardiac fibroblasts

**DOI:** 10.64898/2026.10.20.759862

**Authors:** Huanzhu Jiang, Ganesh Malayath, Austin Kellogg, Nongmaithem Debeni Devi, Charlotte Chen, Nathaniel Huebsch

## Abstract

Human induced pluripotent stem cell-derived cardiac fibroblasts (iPSC-cFBs) offer a renewable, genetically defined stromal cell source for studying the biology of cardiac fibrosis and of cardiomyocyte-fibroblast crosstalk in the context of development and disease. As cardiac fibroblasts are proliferative cells, the ability to expand iPSC-cFB in culture is attractive given the need for large cell numbers for experiments such as drug screening and tissue engineering efforts. However, potential phenotypic drift during expansion – in particular, the prospect of a premature Fibroblast-to-Myofibroblast-Transition (FMT) is a concern. Here, we investigated how differentiation protocol, mechanical culture history, and serial expansion shape iPSC-cFB phenotype, transforming growth factor-β1 (TGF-β1) responsiveness, and fibroblast-autonomous remodeling in three-dimensional (3D) tissues. Initial studies of iPSC-cFBs expanded on conventional tissue-culture polystyrene (TCP) revealed passage-associated increases in projected cell area and expression of α-smooth muscle actin (α-SMA) two markers of FMT. These findings motivated us to study how the rigidity of a poly (dimethyl siloxane) (PDMS; either 15 kPa or 2 MPa) substrate used during initial iPSC-cFB derivation and subsequent expansion impacts both basal phenotype and response to the FMT-inducing growth factor TGF-β. Across iPSC-cFB derived with two independent differentiation protocols, mechanical conditioning produced passage-dependent basal phenotypes and responsiveness to TGF-β. Interestingly, the impact of mechanical pre-conditioning on collective response of iPSC-cFB to TGF-β within “*in vivo*-like” 3D cultures was distinct from the impact of this conditioning when the cells were plated to 2D substrates. Together, these findings demonstrate that mechanical culture history shapes iPSC-cFB response capacity and tissue-remodeling behavior without irreversibly fixing fibroblast state. Passage and subsequent microenvironment can attenuate or re-express these history-dependent phenotypes. This study establishes a foundation for using iPSC-cFB for efforts in basic biological discovery and future tissue engineering efforts.

## INTRODUCTION

Cardiac fibroblasts regulate extracellular-matrix (ECM) homeostasis, myocardial structural integrity, and tissue repair^1,2^. Through ECM remodeling and paracrine signaling, fibroblasts also influence myocardial mechanical and electrical behavior^2,3^. In the heart, fibroblast-driven scarring can diminish contractility and increase risks of arrhythmia. Fibroblast-to-Myofibroblast Transition (FMT) can be induced through both chemical cues (e.g. transforming growth factor-β, TGF-β) and mechanical cues, including matrix rigidity, mechanical stretch, and tensional anisotropy^4–6^. Increased cell size and α-smooth muscle actin (α-SMA) expression are two key markers of the FMT^1,2,7^.

Although primary human cardiac fibroblasts are valuable for studying myocardial remodeling, donor variability and limited scalability can complicate genetically control studies^2,8,9^. Furthermore, in the modern era of “omics,” donor consent and institutional oversight may restrict secondary specimen use or subsequent genomic and transcriptomic data sharing^10^. Human iPSC-cFBs provide an expandable alternative, with differentiation strategies through epicardial or second-heart-field intermediates^8,9,11^. Although they may not fully recapitulate mature adult fibroblasts, their controlled genetic background and scalability make them valuable for mechanistic studies and tissue engineering. Zhang et al. developed an epicardial differentiation protocol yielding relatively quiescent, TGF-β-responsive fibroblasts^9^, which Whitehead et al. subsequently refined to improve fibroblast generation and phenotypic stability^8^. However, differentiation methodology can influence the resulting cellular phenotype. Floy et al. demonstrated that epicardial- and second-heart-field-derived fibroblasts differ in ECM composition, paracrine signaling, and activation potential^12^, while Fernandes et al. demonstrated the emergence of heterogeneous fibroblast populations from epicardial progenitors in cardiac organoids^13^. These findings suggest that cell-derivation history may affect subsequent function.

Culture conditions further alter iPSC-cFB phenotype. Kerr et al. demonstrated that cardiac ECM can attenuate activation-associated characteristics that persist after transfer to a different substrate^14^. Nagalingam et al. further demonstrated passage-associated transcriptional and metabolic remodeling^15^, while Hall et al. showed that chronic fibroblast activation can reduce phenotypic reversibility^16^. Mechanical and biochemical signaling are also closely linked: cyclic stretch can attenuate TGF-β-induced activation, whereas matrix stiffness promotes activation through mechanosensitive pathways^4,17^. Studies using reversibly tunable substrates have additionally demonstrated mechanical memory in cardiac fibroblasts^4^. Nevertheless, how differentiation protocol, mechanical culture history, and serial expansion jointly influence subsequent fibroblast responsiveness and function remains incompletely understood.

Understanding on how iPSC-cFB derivation protocol and subsequent conditions for cellular expansion is particularly relevant to using iPSC-cFB in tissue-engineered models. For example, incorporating iPSC-cFBs into engineered cardiac tissues has been shown to improve calcium handling, tissue compaction, structural organization, and contractile function^12,18,19^. The scalability of producing iPSC-cFB has also enabled high-throughput antifibrotic drug screening and subsequent validation in engineered tissues and preclinical models^20^.

Here, we investigated MYBPC3fs iPSC-cFBs generated using two adapted epicardial protocols and maintained on soft (15 kPa) or rigid (2 MPa) PDMS substrates through passage 2 (P2) or passage 5 (P5). We assessed basal phenotype, subsequent TGF-β responsiveness following transfer to a common 2D environment, and fibroblast-autonomous remodeling in 3D tissues. We hypothesized that differentiation protocol, mechanical culture history, and serial expansion jointly shape subsequent fibroblast behavior, establishing culture history as an important variable in engineered cardiac models.

## Results

### Phenotypic drift of passaged iPSC-cFB

We first examined iPSC-cFBs generated using the Zhang epicardial differentiation protocol^9^ (**Supplemental Fig. 1A**), during expansion on conventional TCP substrates. Consistent with data from these authors, iPSC-cFB derived with this protocol exhibited signs of premature FMT after 5 passages: brightfield imaging showed a more spread morphology, while immunostaining revealed more prominent α-SMA-positive structures from P2 to P5 **(Fig. 1B-E, Supplemental Fig. 1B)**. Surprisingly, even at P2, iPSC-cFB derived from this protocol did not exhibit evidence for FMT when treated with a high dose (10ng/mL) of TGF-β1, a classic chemical profibrotic stimulus **(Fig. 1F-G)**. We verified basal iPSC-cFB phenotypes in a second isogenic control line, harboring a cardiomyopathy-associated frameshift variant in Myosin Binding Protein C without detectable changes in cell area, α-SMA intensity, or eccentricity (*MYBPC3*; **Supplemental Fig. 1C-F**). As MYBPC3 expression is restricted to cardiomyocytes and absent in cardiac fibroblasts, we performed subsequent experiments in the *MYBPC3* frameshift line.

**Figure 1.**
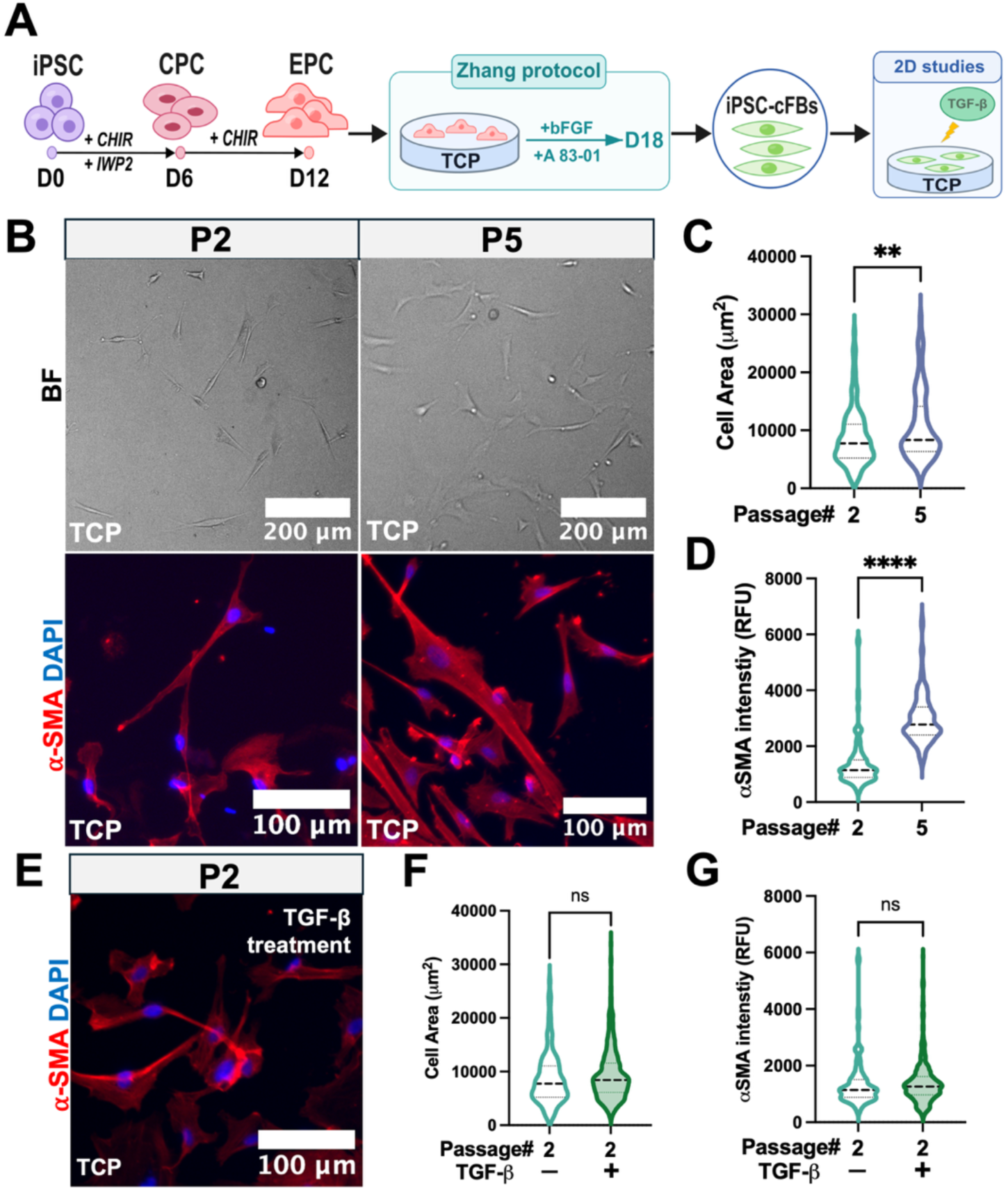
Conventional culture on TCP reveals passage-associated phenotypic drift in iPSC-cFB and motivates a mechanically defined culture system. **(A)** Schematic overview of iPSC-cFB generation using the Zhang differentiation protocol and subsequent experiments under defined mechanical conditions. Following differentiation through cardiac progenitor cell (CPC) and epicardial progenitor cell (EPC) stages, cells were maintained on soft 15kPa or stiff 2MPa PDMS substrates and evaluated at passage 2 (P2) and passage 5 (P5) in 2D cultures and fibroblast-only 3D tissue constructs, with or without TGF-β stimulation. **(B)** Representative brightfield (top) and α-SMA immunostaining (red with blue nuclear counterstain) images of iPSC-cFBs cultured on TCP at P2 and P5. **(C)** Quantification of projected cell area in P2 and P5 iPSC-cFBs maintained on TCP. **(D)** Quantification of α-SMA fluorescence intensity in P2 and P5 isogenic-control iPSC-cFBs maintained on TCP. **(E)** Representative α-SMA immunostaining of P2 iPSC-cFBs cultured on TCP following 10 ng/mL TGF-β1 treatment. **(F, G)** Quantification of **(F)** cell area and **(G)** α-SMA fluorescence intensity in the absence or presence of TGF-β at P2. n values represented ≥ 3 independent batches. Error bar: *SD*. **, **** indicate *p* value less than 0.01 and 0.0001, respectively.

### Substrate stiffness affects passage-dependent phenotypic drift in iPSC-cFB

The challenge of premature FMT in cultured fibroblasts is well-documented, and prior work suggests that this can be suppressed to some extent in primary fibroblasts by passaging the cells on substrates with physiological stiffness^21–23^. We confirmed this phenomenon in human primary cFB by passaging these primary cells 2 times on either PDMS substrates with physiological (∼15 kPa) stiffness^24,25^, or TCP, and then replating to common TCP substrates for subsequent exposure to TGF-β. TGF-β did not significantly alter cell area or α-SMA intensity following TCP conditioning. After two passages on 15 kPa PDMS, TGF-β significantly increased α-SMA fluorescence intensity without a detectable change in cell area **(Supplemental Fig. 2)**.

Based on these findings, we hypothesized that we could prevent premature FMT in passaged iPSC-cFB by expanding these cells on substrates with physiological stiffness (15 kPa). To isolate mechanical effects from substrate surface chemistry, we maintained the cells on PDMS substrates with elastic moduli of either 15 kPa or 2 MPa. Prior work by Palchesko suggests negligible differences in both surface chemistry and surface texture across these conditions^25^. The 2 MPa substrate was used as a deliberately rigid culture condition, well above the stiffness of even fibrotic tissues. To determine whether the method of differentiation might also affect passage and stiffness-dependent FMT, we performed our studies in iPSC-cFB derived from both the original Zhang protocol^9^ and a more recent protocol from Whitehead *et al*^8^., in which additional time is spent in the EPC stage **(Fig. 2A)**. Cells were plated to the PDMS substrates as epicardial progenitor cells (EPC); they completed their transition from EPC to iPSC-cFB on PDMS, and stayed on PDMS substrates for subsequent expansion to either P2 or P5 **(Fig. 2B)**. We compared cells at P2 and P5 (rather than freshly derived P0 iPSC-cFB) to reflect the eventual need to produce iPSC-cFB *en masse* for drug screening and disease modeling applications. In both cases, following expansion on PDMS, we plated iPSC-cFB to common TCP substrates to assess morphology and TGF-β1 response.

**Figure 2.**
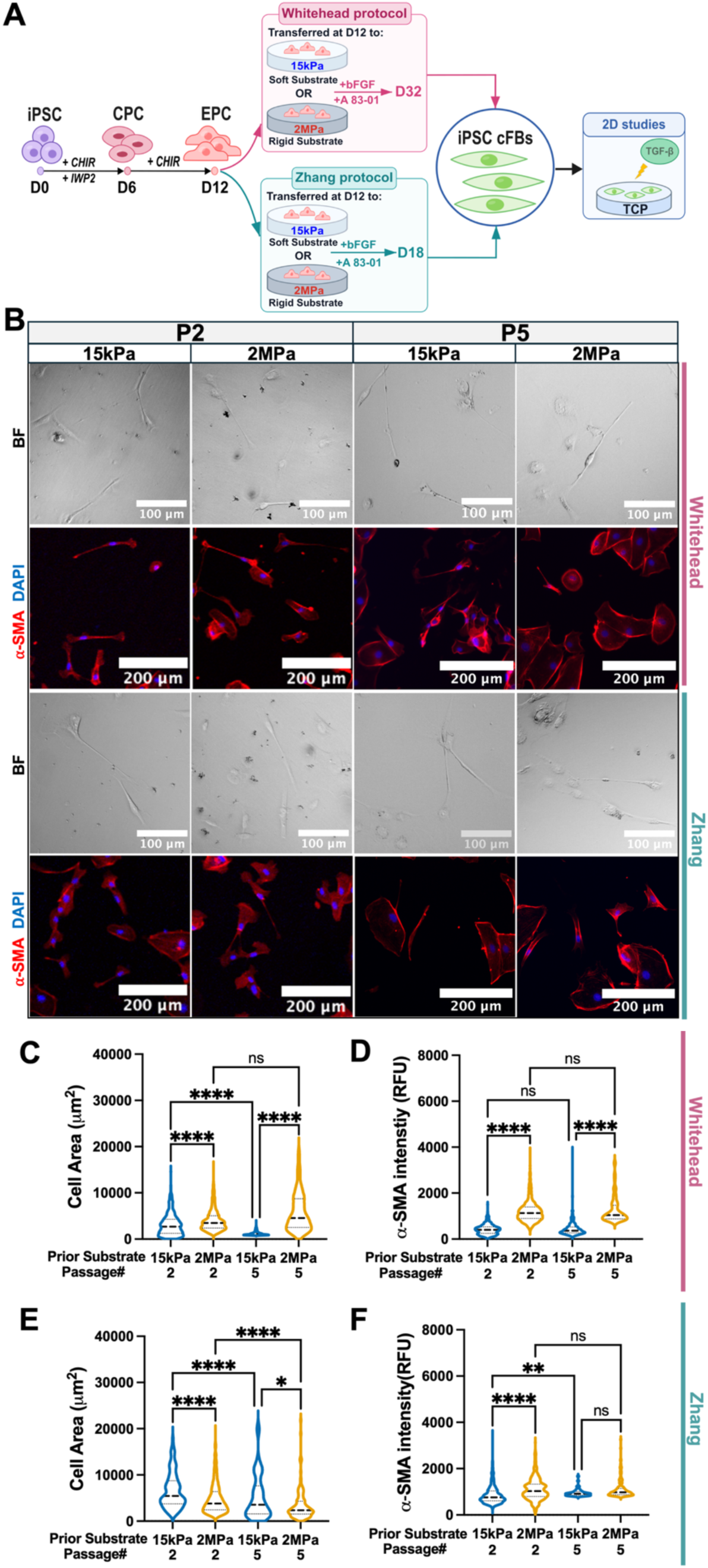
Defined mechanical culture conditions regulate the basal phenotype of iPSC-derived cardiac fibroblasts during serial expansion. **(A)** Representative brightfield and α-SMA immunostaining images of Whitehead-and Zhang-derived iPSC-cFBs at P2 and P5 following culture on 15kPa or 2MPa PDMS substrates. Columns correspond to P2 15 kPa, P2 2MPa, P5 15kPa, and P5 2MPa; **(B)** brightfield and α-SMA/nuclei images are shown for each differentiation protocol. **(C-D)** Quantification of **(C)** cell area and **(D)** α-SMA fluorescence intensity in Whitehead-derived iPSC-cFBs with the indicated mechanical culture histories at P2 and P5. **(E-F)** Quantification of **(E)** cell area and **(F)** α-SMA fluorescence intensity in Zhang-derived iPSC-cFBs with the indicated mechanical culture histories at P2 and P5. n values represented ≥ 3 independent batches. Error bar: *SD*. **** indicate *p* value less than 0.0001.

Despite being plated to a common TCP substrate following expansion on PDMS, Whitehead-protocol derived iPSC-cFB exhibited pronounced differences in morphology and α-SMA expression based on the PDMS substrate used for their expansion. At P2, cells maintained on 2 MPa PDMS had significantly greater projected area than cells that had been expanded on 15 kPa PDMS **(Fig. 2C)**. This difference became more pronounced at P5, when cell area decreased in the soft group but increased in the rigid group. α-SMA fluorescence intensity was also significantly higher on 2 MPa PDMS at both passages **(Fig. 2D)**. In contrast, Zhang-derived cells exhibited greater projected area following 15 kPa conditioning at both passages **(Fig. 2E)**. Nevertheless, α-SMA fluorescence intensity was higher after 2 MPa conditioning at P2, despite their smaller projected area **(Fig. 2F)**. By P5, the α-SMA difference between mechanical conditions was no longer significant, although the difference in cell area persisted **(Fig. 2E-F)**. These findings reveal distinct relationships among cell spreading, α-SMA expression, and serial expansion. In particular, greater cell spreading did not consistently correspond to higher α-SMA expression, suggesting that these measurements capture different aspects of the fibroblast phenotype. It is also possible that contact inhibition masked fibroblast spreading in these studies.

### Prior mechanical culture history and serial expansion influence subsequent TGF-β responsiveness

Having established that mechanical conditioning alters basal iPSC-cFB phenotype, we next asked to determine whether mechanical culture history altered the capacity of iPSC-cFBs to exhibit TGF-β-driven FMT. After expansion on PDMS, cells were replated to common TCP substrates and exposed to 10 ng/mL TGF-β for 48 hr **(Fig. 3A)**. In Whitehead-derived cells, TGF-β increased cell area and α-SMA intensity at P2 following either mechanical history **(Fig. 3B-D)**. However, the response trajectories diverged at P5. 15 kPa-conditioned cells retained significant increases in both measurements, whereas 2 MPa-conditioned cells exhibited reduced cell area and no detectable additional α-SMA induction **(Fig. 3C-D)**. Relative-response analyses further demonstrated greater TGF-β responsiveness following 15 kPa conditioning, particularly at P5 **(Supplemental Fig. 4A-B)**.

**Figure 3.**
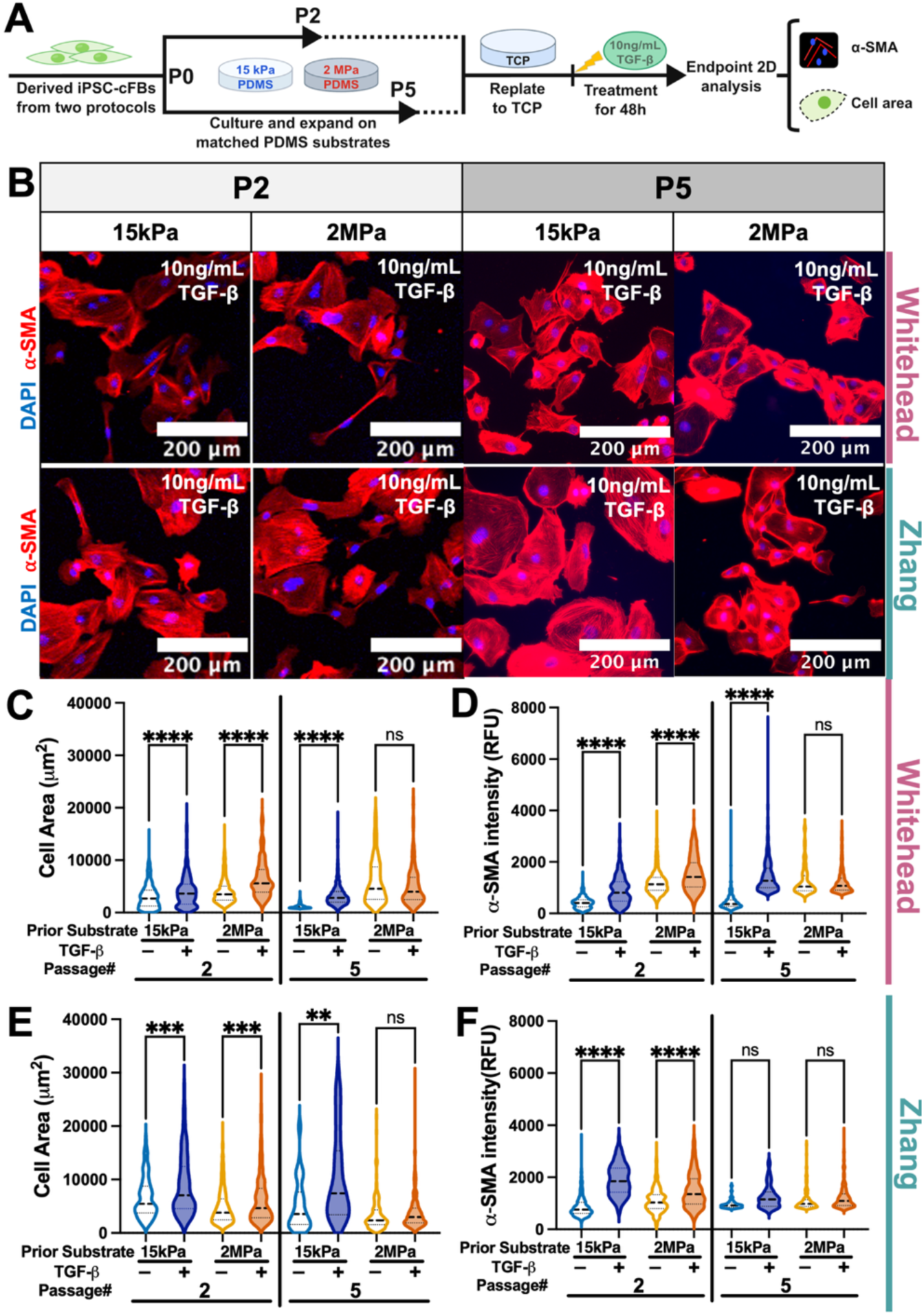
Mechanical culture history modulates subsequent profibrotic responsiveness of iPSC-cFBs. **(A)** Experimental schematic for 2D studies of MYBPC3fs iPSC-cFBs generated using the Zhang or Whitehead differentiation protocol through P2 or P5. **(B)** Representative α-SMA immunostaining of Whitehead- and Zhang-derived iPSC-cFBs following no treatment or TGF-β treatment across passage and prior mechanical-culture conditions. **(C)** Cell area and **(D)** α-SMA fluorescence intensity following no treatment or TGF-β treatment under 15kPa and 2MPa mechanical culture conditions in Whitehead-derived iPSC-cFBs at P2 and P5. **(E)** Cell area and **(F)** α-SMA fluorescence intensity following no treatment or TGF-β treatment under 15kPa and 2MPa mechanical culture conditions in Zhang-derived iPSC-cFBs at P2 and P5. n values represented ≥ 3 independent batches. Error bar: *SD*. *, **** indicate *p* value less than 0.05 and 0.0001, respectively.

Zhang-derived cells exhibited TGF-β-induced cell area increase at both passages following either mechanical history **(Fig. 3E-F)**. In contrast, α-SMA induction was significant at P2 but not P5, with a greater proportional P2 response following 15 kPa conditioning **(Fig. 3F, Supplemental Fig. 4C-D)**. Thus, unlike Whitehead-derived fibroblasts, which retained a strong α-SMA response at P5 following soft conditioning, Zhang-derived fibroblasts exhibited attenuated short-term α-SMA inducibility at P5 following both conditioning environments.

To further characterize treatment-associated morphological remodeling, we quantified cell eccentricity, a measure of projected cell elongation ranging from 0 for circular profiles to near 1 for highly elongated profiles **(Supplemental Fig. 3A)**. TGF-β reduced eccentricity in Whitehead-derived P2 cells following both mechanical histories and in P5 cells following soft conditioning, whereas no significant change was detected in P5 rigid-conditioned cells **(Supplemental Fig. 3B)**. Zhang-derived cells showed reduced eccentricity at P2 following soft conditioning and at P5 following both mechanical histories, but no significant change was detected in P2 rigid-conditioned cells **(Supplemental Fig. 3C)**. These results indicate that treatment-associated increases in projected area frequently coincided with a more rounded, uniformly spread cell shape. Overall, despite protocol-specific differences in response trajectories, prior culture on the 15 kPa substrate was generally associated with greater preservation of TGF-β responsiveness during serial expansion.

### Prior mechanical history produces passage-dependent carryover into fibroblast-only 3D tissue remodeling and mechanics

To determine whether prior mechanical conditioning also influences collective fibroblast behavior within an “*in vivo*-like” 3D context, P2 and P5 iPSC-cFBs were incorporated into engineered tissues by encapsulating the cells into collagen I gel in microwells embedded with two PDMS posts^26–28^ **(Fig. 4A)**. Besides allowing us to assess how passaging history affected collective fibroblast behavior, this approach also allowed us to examine whether differences acquired during 2D expansion persisted after cells were transferred into a common 3D environment. Representative brightfield images and condition-averaged tissue outlines revealed differences in tissue morphology associated with prior mechanical history, particularly at P2 **(Fig. 4A-B)**. At P2, cells previously conditioned on rigid PDMS generated more significant narrower and smaller tissues than soft conditioned cells in both differentiation protocols **(Fig. 4B-D, Supplemental Fig. 5A, E)**. The consistency of this response across the two protocols was notable because Whitehead- and Zhang-derived fibroblasts exhibited differential spreading on TCP depending on their mechanical conditioning during serial passaging **(Fig. 2)**.

**Figure 4.**
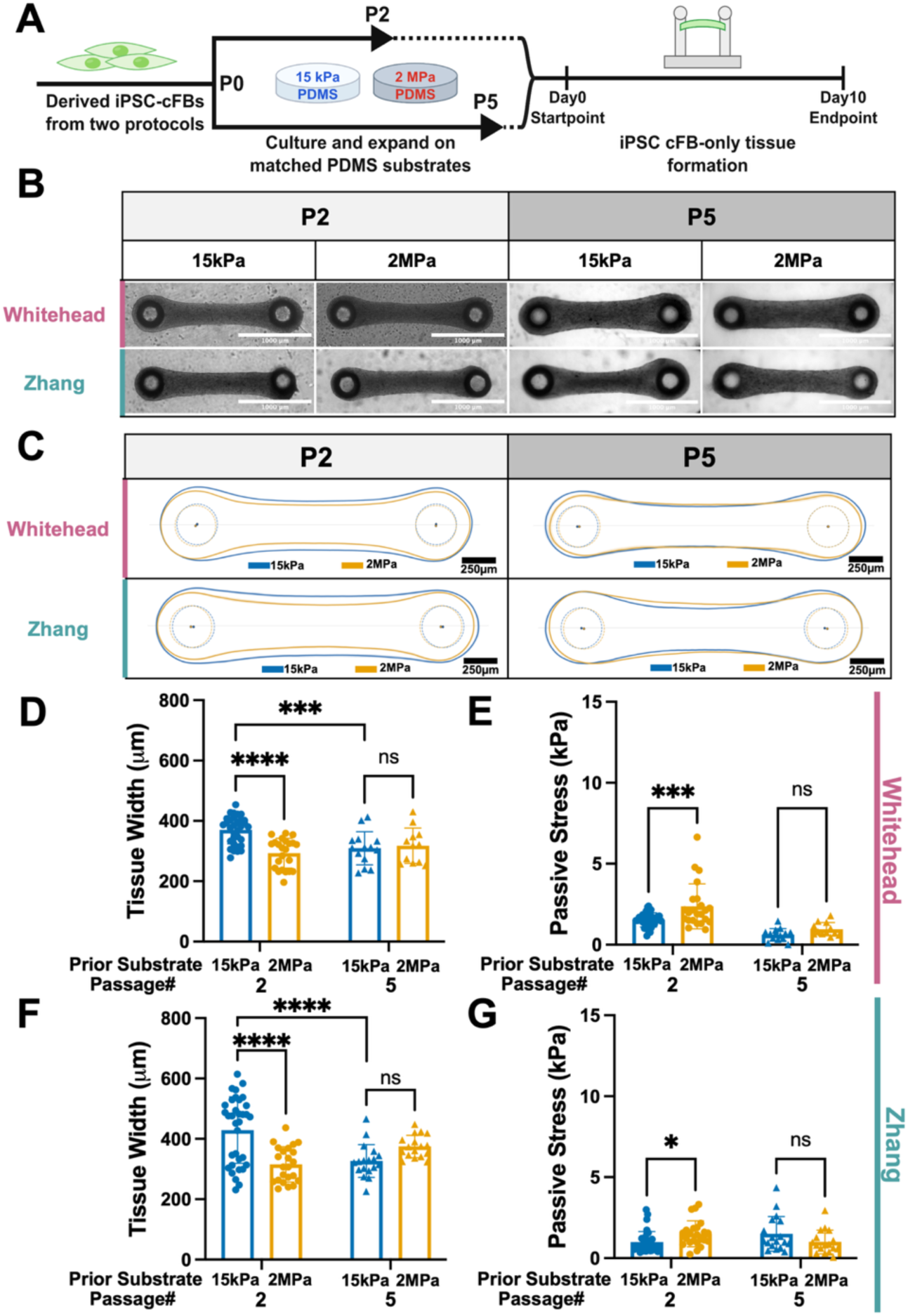
Prior mechanical culture history produces passage-dependent carryover into fibroblast-only 3D tissue remodeling and resting mechanics. **(A)** Experimental schematic for 3D studies of MYBPC3fs iPSC-cFBs generated using the Zhang or Whitehead differentiation protocol through P2 or P5. **(B)** Representative brightfield images of tissues generated from Whitehead- and Zhang-derived iPSC-cFBs with the indicated passage and prior mechanical-culture histories. **(C)** Mean tissue outlines derived from brightfield tissue images across two protocols, passage and prior mechanical culture conditions. Solid contours in tissue outlines represent the condition-averaged tissue outline, and [shaded regions/flanking contours] indicate ± SEM across individual tissue outlines. Dashed circles denote the mean post boundaries, with filled points indicating post centers. **(D-E)** Quantification of endpoint **(D)** longitudinal tissue-width, and **(E)** stress of tissues generated from Whitehead-derived iPSC-cFBs with the indicated mechanical culture histories at P2 and P5. **(F-H)** Quantification of endpoint **(F)** longitudinal tissue-width, and **(G)** stress of tissues generated from Zhang-derived iPSC-cFBs with the indicated mechanical culture histories at P2 and P5. n values represented ≥ 3 independent batches. Error bar: *SD*. *, **, ***, **** indicate *p* value less than 0.05, 0.01, 0.001 and 0.0001, respectively.

Despite these geometric differences, passive tension did not differ significantly between mechanical-history groups at P2 **(Supplemental Fig. 5B, F).** However, passive stress was higher in rigid-history tissues from both protocols **(Fig. 4E, H)**. Because these tissues were narrower and smaller, the elevated normalized stress occurred without a corresponding increase in passive load, indicating that prior mechanical history primarily altered tissue geometry and its relationship to transmitted tension rather than uniformly increasing force generation.

With serial expansion, the magnitude of passive tension decreased from P2 to P5, particularly in Whitehead-derived tissues, while differences between prior 15 kPa and 2 MPa conditions remained nonsignificant within each passage **(Supplemental Fig. 5B, F)**. At the same time, the influence of prior mechanical conditioning was less pronounced at P5. Differences in tissue width and projected size were substantially attenuated, or no longer detectable, and estimated passive stress no longer differed significantly between mechanical-history conditions in either protocol **(Fig. 4C, D, F; Supplemental Fig. 5A, E)**. These findings indicate that the association between prior substrate stiffness and subsequent 3D tissue geometry is strongest at early passage and becomes attenuated during serial expansion.

### TGF-β robustly promotes 3D tissue compaction but has limited effects on passive mechanics

We next investigated whether the stiffness of the 2D substrate used for iPSC-cFB expansion would affect the TGF-β response of 3D iPSC-cFB micro-tissues. Fibroblast-only tissues generated from each mechanical history through P2 or P5 were treated with 10 ng/mL TGF-β from day 2 through day 10 **(Fig. 5A)**. Treatment significantly reduced tissue width across both differentiation protocols, passages, and mechanical history **(Fig. 5B, C, E)**, with corresponding reductions in tissue size **(Supplemental Fig. 5C, G)**. Thus, despite substantial differences in basal phenotype and short-term 2D responsiveness, all populations retained the capacity to undergo pronounced tissue-level compaction following TGF-β exposure.

**Figure 5.**
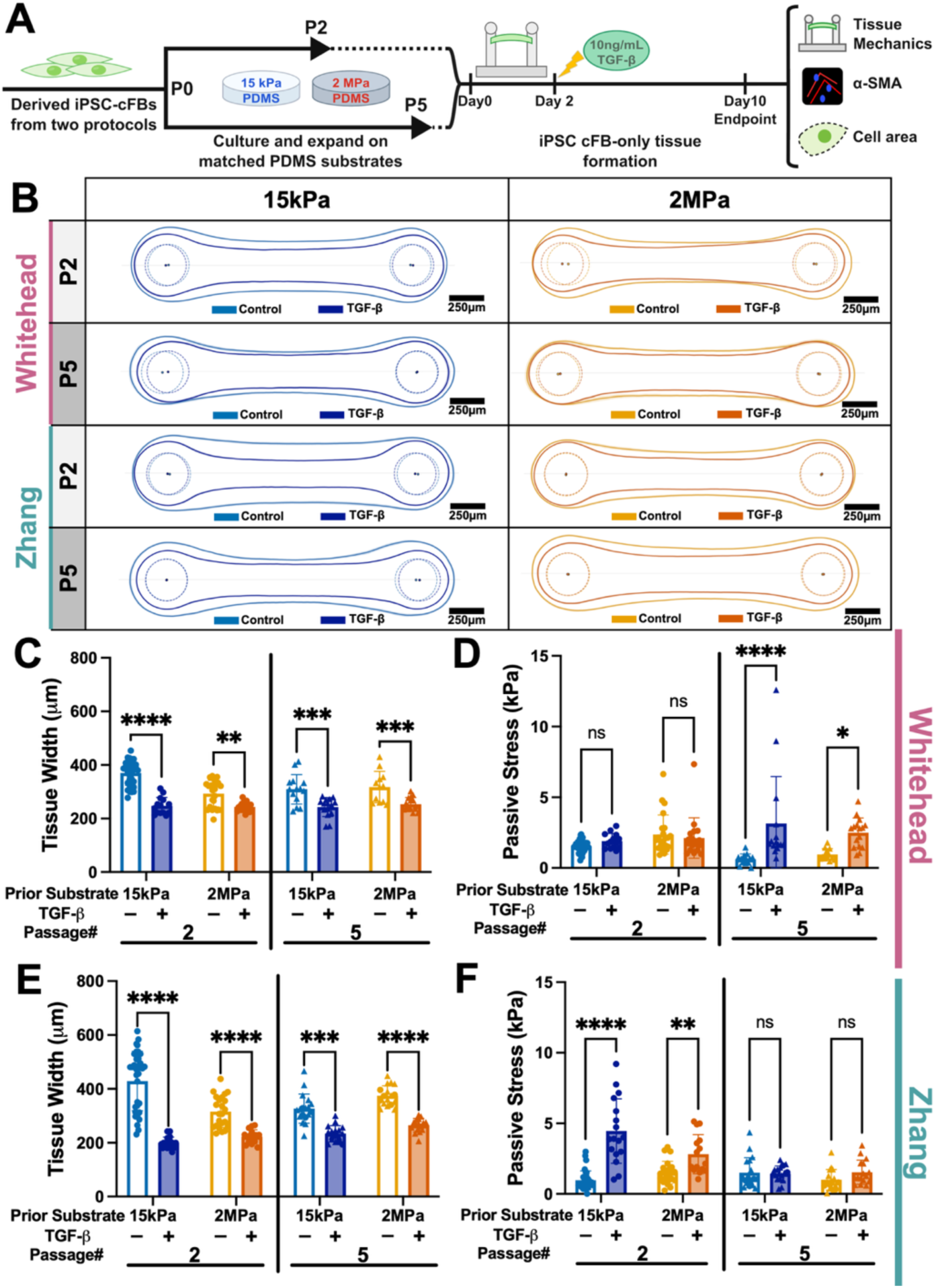
TGF-β alters the morphology and mechanical properties of iPSC-cFB-only tissues following compliant mechanical culture. **(A)** Schematic of 3D experimental design showing MYBPC3fs iPSC-cFBs maintained on 15kPa or 2MPa PDMS through P2 or P5, harvested, and subsequently incorporated into identical fibroblast-only two-post tissue constructs. **(B)** Condition-averaged tissue outlines following no treatment or TGF-β1 treatment across differentiation protocol, passage, and prior mechanical-history conditions. Solid contours represent the mean tissue boundary, with [flanking contours/shaded regions] indicating ± SEM across individual tissue outlines. Dashed circles denote the mean post boundaries, and filled points indicate post centers. **(C-D)** Quantification of endpoint **(C)** tissue width and **(D)** stress for Whitehead-derived tissues at P2 and P5 following no treatment or TGF-β treatment. **(E-F)** Quantification of endpoint **(E)** tissue width and **(F)** stress for Zhang-derived tissues at P2 and P5 following no treatment or TGF-β treatment. n values represented ≥ 3 independent batches. Error bar: *SD*. **, **** indicate *p* value less than 0.01 and 0.0001, respectively.

In contrast, passive mechanical responses highly depend on passage, derivation protocol, and prior mechanical conditioning **(Supplemental Fig. 5D, H)**. In Whitehead-derived tissues, treatment significantly reduced passive tension following both soft and rigid conditioning. By P5, TGF-β reduced passive tension at P2 following both conditioning histories, whereas at P5 it increased tension following soft conditioning but produced no significant change after rigid conditioning **(Supplemental Fig. 5D)**. In Zhang-derived tissues, passive tension was unchanged at P2 and decreased only in soft-history tissues at P5 **(Supplemental Fig. 5H)**.

Estimated passive stress exhibited a distinct pattern **(Fig. 5D, F)**, largely driven by changes in tissue geometry (e.g. by reducing tissue width to reduce cross-sectional area). In Whitehead-derived tissues, TGF-β increased stress at P5, but not P2, following either mechanical history **(Fig. 5D)**. Conversely, Zhang-derived tissues exhibited significant stress increase at P2, but not P5 **(Fig. 5F)**.

### Prior mechanical history remains detectable in the cellular activation-associated phenotype within 3D tissues

To determine whether tissue-level responses were accompanied by cellular responses, we quantified cell area and α-SMA intensity following TGF-β treatment **(Fig. 6A, Supplemental Fig. S6)**. TGF-β significantly increased α-SMA intensity across both differentiation protocols, passages and mechanical histories (**Fig. 6C, E**). Cell-area responses were more variable: significant increases were observed in Whitehaed-derived tissues at P2 following soft conditioning and at P5 following both histories, whereas Zhang-derived tissues exhibited significant increases only at P5 **(Fig. 6B, D)**.

**Figure 6.**
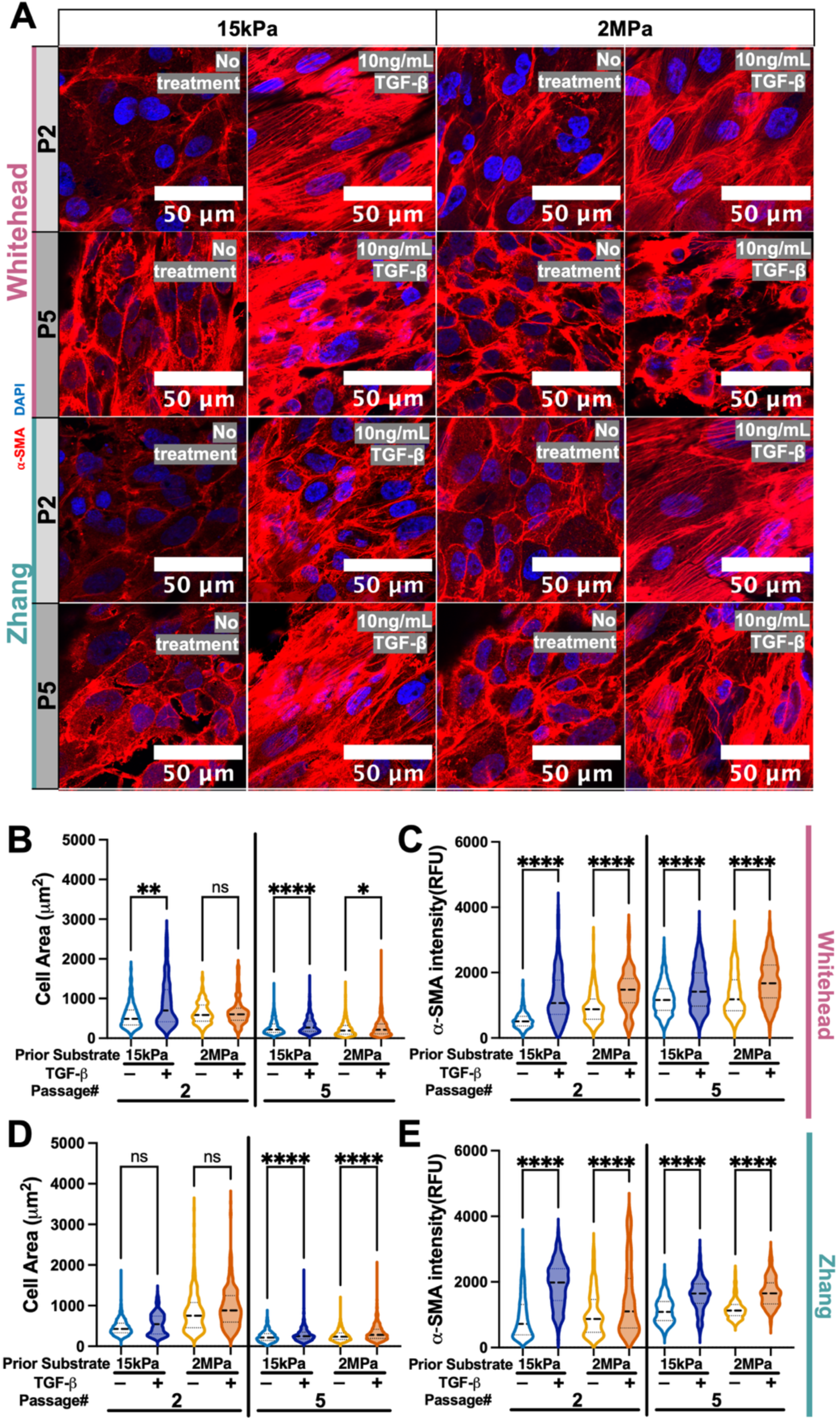
Mechanical culture history and TGF-β1 jointly shape the activation-associated phenotype of iPSC-cFBs within fibroblast-only 3D tissues. **(A)** Representative α-SMA immunostaining (red with blue nuclear counterstain of micro-tissues generated from iPSC-cFB derived via the Whitehead and Zhang protocols that were subsequently expanded on either 15kPa or 2MPa PDMS substrates. A subset of micro-tissues were exposed to 10 ng/mL TGF-β from days 2-10. Images are shown across P2 and P5 conditions. **(B-C)** Quantification of **(B)**cell area and **(C)** α-SMA fluorescence intensity within tissues generated from Whitehead-derived iPSC-cFBs tissues across passage, prior mechanical-history, and TGF-β treatment conditions. **(D-E)** Quantification of **(D)**cell area and **(E)** α-SMA fluorescence intensity within tissues generated from Zhang-derived iPSC-cFBs tissues across passage, prior mechanical-history, and TGF-β treatment conditions. n values represented ≥ 3 independent batches. Error bar: *SD*. *, **, ***, **** indicate *p* value less than 0.05, 0.01, 0.001 and 0.0001, respectively.

Notably, Whitehead-derived P5 rigid-conditioned cells and Zhang-derived P5 cells from both histories exhibited significant α-SMA induction in 3D despite limited responses during the 48-hr 2D challenge **(Fig. 3D, F**; **Fig. 6C, E)**. Relative-response analyses further revealed greater TGF-β-induced α-SMA expression following soft conditioning at P2 in both protocols, with no significant history-dependent difference at P5 **(Supplemental Fig. 7B, D)**. Cell-area response ratios showed a similarly pronounced P2 mechanical-history difference in Whitehead-derived tissues, whereas the corresponding Zhang-derived P5 comparison favored rigid conditioning **(Supplemental Fig. 7A, C)**. Together, these findings show that limited short-term responsiveness in 2D does not necessarily indicate irreversible loss of TGF-β responsiveness under subsequent 3D culture conditions.

## Discussion

In this study, we show that iPSC-derived cardiac fibroblast (iPSC-cFB) phenotype (particularly, premature FMT) is strongly shaped by mechanical culture history. Across the dataset, three themes emerge. First, the effects of mechanical conditioning varied with differentiation protocol and passage, indicating that substrate stiffness does not impose a uniform fibroblast phenotype. Second, prior mechanical conditioning influences subsequent TGF-β responsiveness and 3D remodeling even after the conditioning environment is removed, with soft conditioning associated with greater inducible capacity in several key conditions. Third, these history-dependent states are not fixed: early effects of mechanical history on geometry and stress within 3D engineered tissues diminish after serial expansion, while populations with limited short-term responses to TGF-β exposure in 2D culture retain robust responsiveness in 3D. Together, these findings support a model in which mechanical history is an important determinant of fibroblast response capacity and engineered tissue behavior.

One important finding is that differentiation history modifies how iPSC-cFBs respond to the same mechanical environment. Whitehead-derived cells exhibited greater α-SMA expression following rigid conditioning at both passages, whereas Zhang-derived cells showed greater α-SMA under rigid conditions at P2 but not P5. Morphological responses differed further: Whitehead-derived cells spread more extensively on rigid substrates, whereas Zhang-derived cells spread more on soft substrates, despite greater α-SMA under rigid conditions at P2. Thus, neither substrate stiffness nor cell area alone was sufficient to predict the resulting activation-associated α-SMA expression. This may be associated with changes in local cell density within our cultures, as cell density has been shown to have a direct impact on FMT^29^.

Our findings are consistent with previous evidence that the route used to generate iPSC-cFBs can influence their functional properties. Whitehead et al. showed that epicardial differentiation and expansion conditions affect fibroblast morphology, ECM production, and phenotypic stability^8^, while Floy et al. demonstrated that developmental lineage influences ECM composition, secreted factors, and myofibroblast activation potential in hPSC-derived cardiac fibroblasts^12^. Although both protocols examined here proceed through an epicardial intermediate, our data extend these observations by showing that differences established during differentiation and expansion can influence how fibroblasts subsequently respond to chemomechanical cues. Serial expansion further reshaped these relationships. Zhang-derived isogenic-control cells maintained on conventional TCP exhibited increased cell area and α-SMA from P2 to P5. Nagalingam et al. similarly reported progressive transcriptional and metabolic remodeling during serial passage of hiPSC-cFBs, including increased expression of collagen, periostin, fibronectin, and ECM-processing genes^15^. In our mechanically defined cultures, however, passage did not produce a uniform phenotype: stiffness-dependent α-SMA differences persisted in Whitehead-derived cells but diminished in Zhang-derived cells, while cell morphology followed protocol-specific trajectories. Passage therefore appears to interact with the environment in which expansion occurs rather than functioning as an independent, uniformly activating variable.

A second important finding is that mechanical conditioning altered the capacity of cells to respond to a subsequent profibrotic stimulus. After transfer to a common TCP environment, Whitehead-derived P5 cells retained robust TGF-β-induced increases in both cell area and α-SMA after soft conditioning, whereas rigid-conditioned cells exhibited reduced cell area and no detectable additional α-SMA induction. Zhang-derived cells maintained cell-area responsiveness across passages, but their short-terim α-SMA response diminished at P5 following either mechanical history. Thus, soft conditioning preserved greater TGF-β inducibility in specific conditions rather than uniformly maintaining fibroblast quiescence. These findings are consistent with previous studies providing precedent for such history-dependent behavior. Kerr et al. showed that hiPSC-cFBs cultured on cardiac ECM retained reduced expression of selected activation-associated genes after subsequent transfer to TCP^14^, demonstrating that culture-associated phenotypes can persist beyond the original environment. Hall et al. conversely found that prolonged activation of primary human and iPSC-derived cFB reduced the reversibility of the myofibroblast phenotype after transfer to soft substrates or inhibition of TGF-β receptor signaling^16^. Bouhrira et al. more directly demonstrated mechanical memory in cardiac fibroblasts using reversibly tunable substrates, showing that prolonged stiff priming influenced subsequent cell area and α-SMA responses after the mechanical environment was changed^30^. Together, these studies support the broader concept that fibroblasts integrate prior mechanical experience rather than responding exclusively to their immediate environment.

Recent work also provides potential mechanistic context. Cho et al. demonstrated stiffness-dependent activation of human iPSC-cFBs and identified SRC-associated matrix mechanosensing as an important component of persistent fibroblast activation^4^. However, the present experiments do not identify a specific molecular mechanism of mechanical memory. Cell harvesting, replating, deposited ECM, cytoskeletal organization, transcriptional adaptation, or population selection could all contribute to the observed persistence. We therefore interpret our findings as functional carryover of mechanical culture history rather than evidence for a specific memory pathway.

The fibroblast-only 3D tissues demonstrate that this carryover extends beyond isolated-cell phenotype. At P2, rigid-conditioned cells formed narrower and smaller tissues than soft-conditioned cells in both differentiation protocols, despite the contrasting stiffness-dependent cell-area patterns observed in 2D. This indicates that monolayer morphology alone did not directly predict subsequent tissue organization. Rigid-conditioned tissues exhibited greater passive stress without significantly greater passive tension. Because stress is normalized to estimated tissue cross-sectional area, we do note that most of the changes in mechanical stress in tissues were due to differences in tissue geometry rather than robust changes in passive tension. Importantly, this stiffness-history effect on geometry and stress diminished by P5, while Whitehead-derived tissues also exhibited reduced passive tension with serial expansion. These data indicate that prior mechanical history influences the relationship among tissue geometry, remodeling, and transmitted passive load, but their effects evolve with serial expansion. While an ability to passage iPSC-cFB indefinitely would certainly be attractive given the need to produce large numbers of these cells for drug screening and disease modeling applications, these findings argue for more limited expansion of the freshly-derived iPSC-cFB, as premature FMT cannot be suppressed by culturing these cells on soft substrates.

TGF-β treatment revealed substantial retained remodeling capacity across culture histories. Tissue width and overall size decreased consistently across differentiation protocols, passages, and prior stiffness conditions. In contrast, passive tension and stress showed distinct, condition-dependent responses. This separation is consistent with previous evidence that fibroblast-mediated tissue compaction and mechanical properties can be regulated differentially^31^. Importantly, TGF-β increased α-SMA expression under the 3D treatment conditions, including in populations with limited short-term α-SMA induction in 2D. At P2, soft conditioning was also associated with greater proportional α-SMA induction in 3D in both protocols. Thus, diminished responsiveness in a short-term 2D assay did not represent irreversible loss of profibrotic capacity, which perhaps is the strongest evidence for retained fibroblast plasticity.

Defining this plasticity is particularly important for creating tissue-engineered disease models – there, pre-passaging conditions may influence not only basal iPSC-cFB phenotype, but also the interactions the cFB have with other cells in the tissue. For example, in tissue-engineered models of hypertrophic cardiomyopathy (HCM), including ours, cardiomyocytes harboring sarcomeric mutations can alter stromal behavior through cardiomyocyte-derived signals^32^, but cFB can conversely impact cardiomyocytes’ hypertrophic remodeling^27^. Given that FMT is part of the disease process of HCM, controlling the mechanical history of iPSC-cFBs may be critical for distinguishing disease-specific cardiomyocyte–fibroblast interactions from phenotypic differences introduced during cell expansion.

Several limitations should be considered. The 15 kPa and 2 MPa substrates represent two discrete mechanical environments, with 2 MPa serving as a deliberately rigid condition rather than a direct mimic of fibrotic myocardium. While different PDMS blends can achieve fetal-heart-like stiffness of <1 kPa^24^ (which might further suppress FMT), we purposefully avoided the softest substrates because of challenges in working with these materials in flasks used for cell passaging. The Zhang and Whitehead protocols also differ in multiple differentiation parameters, preventing protocol-dependent effects from being attributed to a single mechanism. Finally, the 2D and 3D TGF-β experiments differed in treatment duration, ECM environment, cellular organization, and mechanical constraint, and cell area and α-SMA represent only selected dimensions of fibroblast phenotype.

Establishing these effects provides a necessary foundation for genetically matched cardiomyocyte-fibroblast models designed to distinguish fibroblast-intrinsic, cardiomyocyte-autonomous, and multicellular mechanisms of inherited diseases like HCM. Future studies will build on this platform by directly testing how fibroblast genotype interacts with mechanical history and cardiomyocyte genotype. This may be particularly important in cardiomyopathies related to genomic variants that impact structures outside the sarcomere, as both cardiomyocytes and fibroblasts could exhibit cell-autonomous changes. Finally, targeted perturbation of mechanosensitive and TGF-β-associated pathways, coupled with transcriptomic or epigenetic profiling, could identify mechanisms that encode and remodel culture-history-dependent fibroblast states.

## Materials and Methods

### Culture and maintenance of iPSC lines

Experiments were performed using an *MYBPC3* frameshift (MYBPC3fs) iPSC line and its isogenic-control^27^. Both lines were maintained in StemFlex media (Life Technologies; A3349401) on plates coated with Geltrex (Life Technologies; A1413302) at 18.75 mg/cm^2^. Cells were passaged using Gentle Dissociation Reagent (PBS containing 1.8g/L NaCl and 0.5mM EDTA) and replated as single cells in StemFlex media supplemented with 10 µM Y-27632 (Biogem; 1293823).

The isogenic-control line was used for the initial TCP studies and the P2 genotype-control comparison, whereas subsequent defined-mechanics experiments were performed using MYBPC3fs iPSC-cFBs.

### Generation and maintenance of iPSC cardiac fibroblasts

iPSC-cFBs were generated using epicardial differentiation strategies adapted from Zhang et al. and Whitehead et al^8,9^. Both routes generated cardiac progenitor and epicardial intermediates before fibroblast specification. Briefly, iPSCs were treated with 6 μM CHIR99021 for 48 hr, allowed to recover for 24 h, treated with 5 μM IWR-1 (Millipore Sigma; I0161) for 48 hr, and then allowed to recover for an additional 24 h. On day 6, cardiac progenitor cells were replated at 20,000 cells/cm² on Geltrex-coated TCP in Advanced DMEM (Fisher Scientific; 12491015) containing 2.5 mM GlutaMAX (Fisher Scientific; 35050061) and 60 μg/mL ascorbic acid (AA) (Fisher Scientific; NC0602549) with 5 μM Y27632 supplementation.

From day 6 onward, cells were directed toward the epicardial lineage using protocol-specific conditions. For the Zhang protocol, cells were treated with 5 μM CHIR99021 and 2 μM retinoic acid (Millipore Sigma; R2625) for 48 hr, followed by recovery in basal media (Advanced DMEM/Glutamax/AA) through approximately day 12. For the Whitehead-derived workflow, cells were treated with 3 μM CHIR99021 without retinoic acid during epicardial specification, followed by culture in basal media through day 12. At approximately day 12, epicardial-progenitor cells (EPC from both workflows were transferred to the assigned 15 kPa or 2 MPa PDMS substrate where they completed fibroblast differentiation and were subjected to subsequent expansion under their respective protocol conditions **(Fig. 2A)**. Zhang-derived cells underwent shorter differentiation in fibroblast media consisting of fibroblast growth medium (PromoCell; C-23025), 10 ng/mL bFGF (Fisher Scientific; 100-18B), and TGF-β Receptor inhibition (0.5 µM A83-01; Bio-Techne 2939) through approximately day 18, whereas Whitehead-derived cells underwent prolonged exposure to 10 ng/mL bFGF-supplemented basal media (Advanced DMEM/Glutamax/AA) through approximately day 32. 0.5 µM A83-01 was also incorporated into the adapted Whitehead workflow during fibroblast differentiation.

Cells were dissociated with 0.25% trypsin-EDTA (Fisher Scientific; 25200072) to generate single-cell suspensions. Following differentiation, cells from both protocols were maintained on their assigned PDMS substrates and expanded to P2 or P5 in fibroblast growth medium supplemented with 10 ng/mL bFGF and 0.5 μM A 83-01.

### Culture of primary cardiac fibroblasts

Human primary ventricular cardiac fibroblasts (Lonza; CC-2904) were maintained in fibroblast growth medium consisting of FBM™ Basal Medium (Lonza; CC-3131) supplemented with FGM™-2 SingleQuots™ (Lonza; CC-4126). Cells were dissociated using 0.25% trypsin-EDTA to generate single-cell suspensions. For mechanical-history studies, cells were maintained either on TCP or transferred to 15 kPa PDMS and expanded for two passages under the assigned condition. Following this conditioning period, both groups were replated onto TCP for subsequent experiments.

### PDMS substrate preparation and mechanical conditioning

PDMS substrates with target elastic moduli of 15 kPa and 2 MPa were prepared using a 1:20 blend of Sylgard 184 527 and pure Sylgard 184, respectively. Substrates were cured overnight at 60°C, sterilized by immersion in 70% ethanol followed by overnight UV exposure, and coated with Geltrex at 18.75 mg/cm² using the same procedure for both stiffness conditions. The 15 kPa substrate was designated “soft” and the 2 MPa substrate “rigid.” Cells remained on their assigned substrate through P2 or P5 unless harvested for downstream assays.

### 2D isogenic-control and primary cardiac fibroblast studies

For initial studies under conventional culture conditions, Zhang-derived isogenic-control iPSC-cFBs were maintained on TCP. Basal fibroblast phenotype was compared between P2 and P5 cells. To assess profibrotic responsiveness under conventional culture conditions, P2 cells were replated onto TCP at 3000 cells/cm^2^ and treated with 10 ng/mL TGF-β1 (PeproTech; AF-100-36E-10UG) for 48 hr. A separate P2 genotype-control experiment compared Zhang-derived isogenic control and MYBPC3fs iPSC-cFBs cultured under the same TCP conditions.

For primary cardiac fibroblast mechanical-history studies, human ventricular cardiac fibroblasts were maintained either on TCP or transferred to 15 kPa PDMS and expanded for two passages under the assigned condition. Cells from both groups were then harvested, replated onto TCP at 3000 cells/cm^2^ under a common mechanical environment, and treated with 10 ng/mL TGF-β1 for 48 hr.

### 2D phenotyping and TGF-β treatment

For the primary mechanobiology experiments, MYBPC3fs iPSC-cFBs generated using the Zhang- and Whitehead-derived differentiation workflows were maintained on their assigned 15 kPa or 2 MPa PDMS substrates through P2 or P5. To evaluate whether prior mechanical conditioning influenced subsequent profibrotic responsiveness, P2 and P5 MYBPC3fs iPSC-cFBs from each substrate condition were harvested and replated onto TCP at 3000 cells/cm^2^, thereby placing all groups in a common mechanical environment. Cells were then treated with 10 ng/mL TGF-β1 for 48 hr before fixation and analysis of cell area, cell eccentricity, and α-SMA fluorescence intensity.

### Immunofluorescence and 2D image analysis

Cells were fixed in 4% paraformaldehyde for 20 min and washed three times with dPBS. 2D monolayer cultures were stained for vimentin (Fisher Scientific; MA5-11883) and α-smooth muscle actin (α-SMA; Abcam; ab124964), with Hoechst 33342 used for nuclear counterstaining.

Images were acquired on an epifluorescence microscope (Nikon Eclipse Ts2R; Tokyo, Japan) equipped with a Hamamatsu ORCA-Flash4.0 V2 digital CMOS camera and an Aura II epifluorescence illumination system with 350-, 488-, 540-, and 647-nm LED light sources. All samples within a given experiment were imaged using identical acquisition settings, including a 10-ms exposure time.

Cell area, α-SMA fluorescence intensity and cell eccentricity were quantified from background-corrected images using custom image-processing software developed in-house **(Supplemental Fig. 3A)**.

### Fibroblast-only two-post tissue formation

P2 or P5 iPSC-cFBs were harvested from their assigned 15 kPa or 2 MPa substrates and embedded in an ECM hydrogel consisting of 1.5 mg/ml collagen I rat tail (Fisher Scientific; A1048301) and 10% v/v Geltrex, and the mixture was neutralized with 1M NaOH to physiological pH, at a density of 1×10^7^ cells/mL.

P2 or P5 iPSC-cFBs were harvested from their assigned 15-kPa or 2-MPa substrates and suspended at 1 × 10⁷ cells/mL in an ECM hydrogel containing 1.5 mg/mL rat-tail collagen I (Fisher Scientific; A1048301) and 10% (v/v) Geltrex. The hydrogel was neutralized to physiological pH with 1 M NaOH, and the cell– matrix suspension was cast into two-post tissue devices fabricated from pure Sylgard 184^27^. To isolate the effect of prior mechanical conditioning, cells from both substrate-history groups were incorporated into the same 3D matrix formulation at the same cell density and cultured in devices with identical geometry.

### 3D TGF-β treatment, tissue geometry, and mechanics

For 3D studies, tissues were treated with 10 ng/mL TGF-β beginning on day 2 and maintained through the day 10 endpoint **(Fig. 5A)**. Tissue morphology was quantified from brightfield images. Tissue outlines were extracted, aligned to the two-post geometry, and averaged within each experimental condition. Solid contours represent the mean tissue boundary, flanking contours or shaded regions represent ±SEM across individual tissue outlines, dashed circles indicate mean post boundaries, and filled points denote post centers. Endpoint tissue width and projected tissue size were quantified from the same brightfield images.

Passive tension was calculated from post deflection^27^,

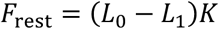

where *K* is calibrated effective post-bending stiffness, *L*_0_ is the unloaded post spacing, and *L*_1_ is the measured post spacing in the presence of the tissue.

Tissue stress was calculated by normalizing passive tension to the estimated tissue cross-sectional area^27^. Cross-sectional area was approximated assuming a circular cross-section,

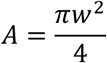

Where *w* is the measured width of an individual tissue. Tissue stress was then calculated as,

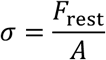

### 3D immunofluorescence and cellular analysis

Following mechanical measurements at the experimental endpoint, tissues were fixed at 4°C using sequential paraformaldehyde concentrations of 1%, 2%, 3%, and 4%, with each step performed for at least 1 h. Fixed tissues were washed three times with dPBS and stained for α-smooth muscle actin (α-SMA; Abcam; ab124964), with Hoechst 33342 used for nuclear counterstaining. Whole tissues were mounted in ProLong Gold antifade reagent (Fisher Scientific; P36930) and imaged using an Olympus Fluoview FV1200 confocal microscope.

Cell area and α-SMA fluorescence intensity within the tissues were quantified using custom image-processing software developed in-house with consistent analysis parameters across experimental conditions **(Supplemental Fig. 6)**.

### Statistical analysis

Statistical analyses and figure generation were performed using GraphPad Prism version 9.3.1. Data are reported as mean ± standard deviation (SD), with experiments conducted using samples from at least three independent batches. Depending on the experimental design, comparisons between two groups were evaluated using the Mann-Whitney U test, while comparisons involving multiple groups were assessed using the Kruskal-Wallis test or two-way ANOVA with Holm-Šidák post-hoc multiple-comparisons testing, as appropriate. All statistical tests were two-sided, and p < 0.05 was considered statistically significant.

## Ethics Approval

Ethics approval is not required.

## Supplemental Materials

See the Supplementary Material for the additional experimental results and supplementary figures, as well as a supporting figure that further validates the findings presented in the main text.

## Acknowledgement

We thank Dr. Michael Vahey (Washington University in St. Louis) for providing access to and assistance with confocal microscopy. This work was supported by the National Heart, Lung, and Blood Institute (R01HL159094, N.H.; R01HL170079, S.C.), the National Science Foundation CAREER Award (2338931), and fellowships from the National Institutes of Health (Austin NIAMS supplement) and American Heart Association to Austin Kellogg.

