## Supplemental Information for "Mechanical history shapes phenotypic plasticity and fibroblast-autonomous 3D remodeling in iPSC-derived cardiac fibroblasts"

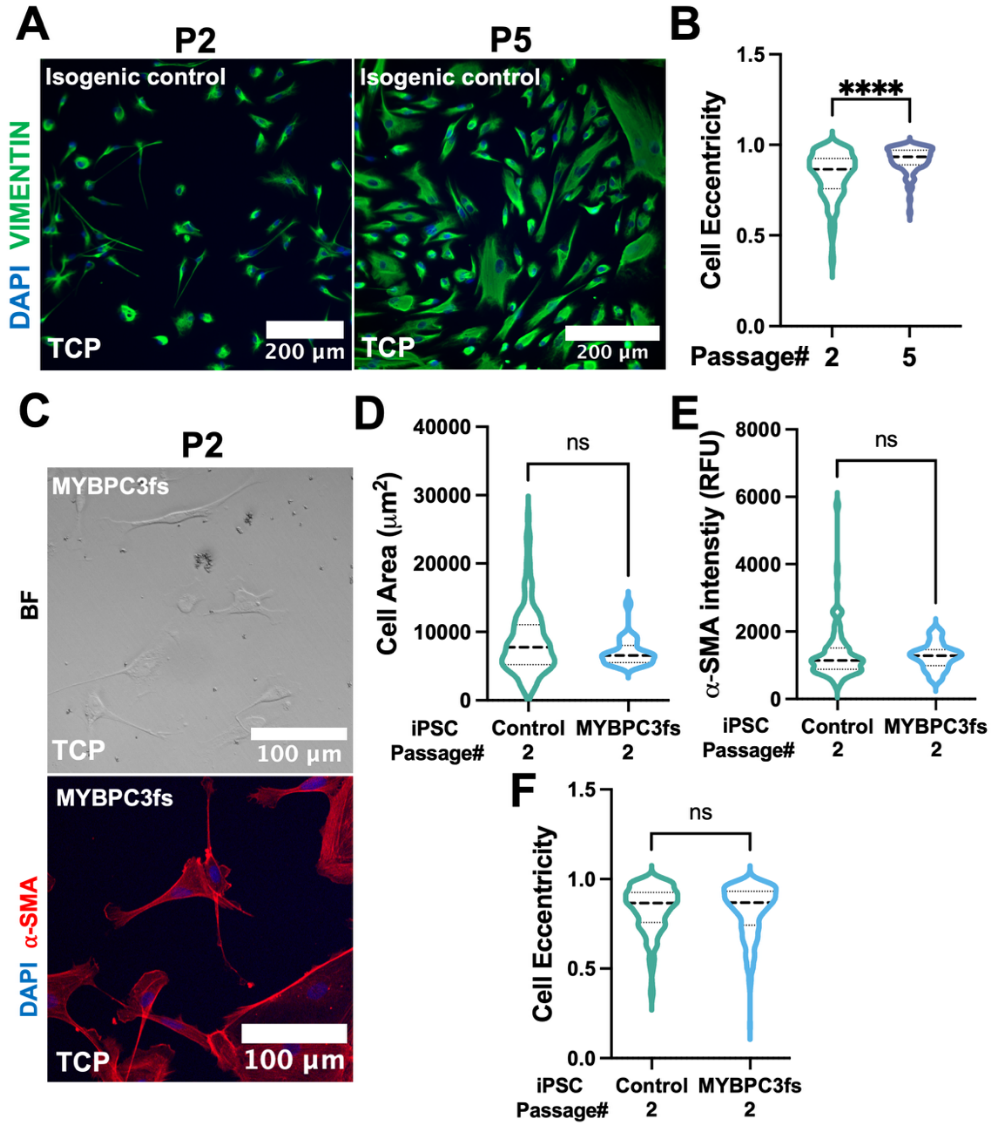


**Figure S1. MYBPC3fs and isogenic-control iPSC-cFBs exhibit comparable basal phenotypes under TCP culture conditions. (A)** Representative Vimentin (green)/nuclei (blue) images of Zhang-derived isogenic-control iPSC-cFBs at P2 and P5 on TCP. **(B)** Quantification of cell eccentricity in Zhang-derived isogenic-control iPSC-cFBs at P2 and P5 on TCP**. (C)** Representative brightfield image and α-SMA (red)/nuclei (blue) images of Zhang protocol-derived MYBPC3fs iPSC-cFBs cultured on TCP at P2. **(D-F)** Comparison of **(D)** cell area, **(E)** α-SMA fluorescence intensity, and **(F)** cell eccentricity between Zhang-derived isogenic-control and MYBPC3fs iPSC-cFBs at P2 under TCP culture conditions. Error bar: *SD*.

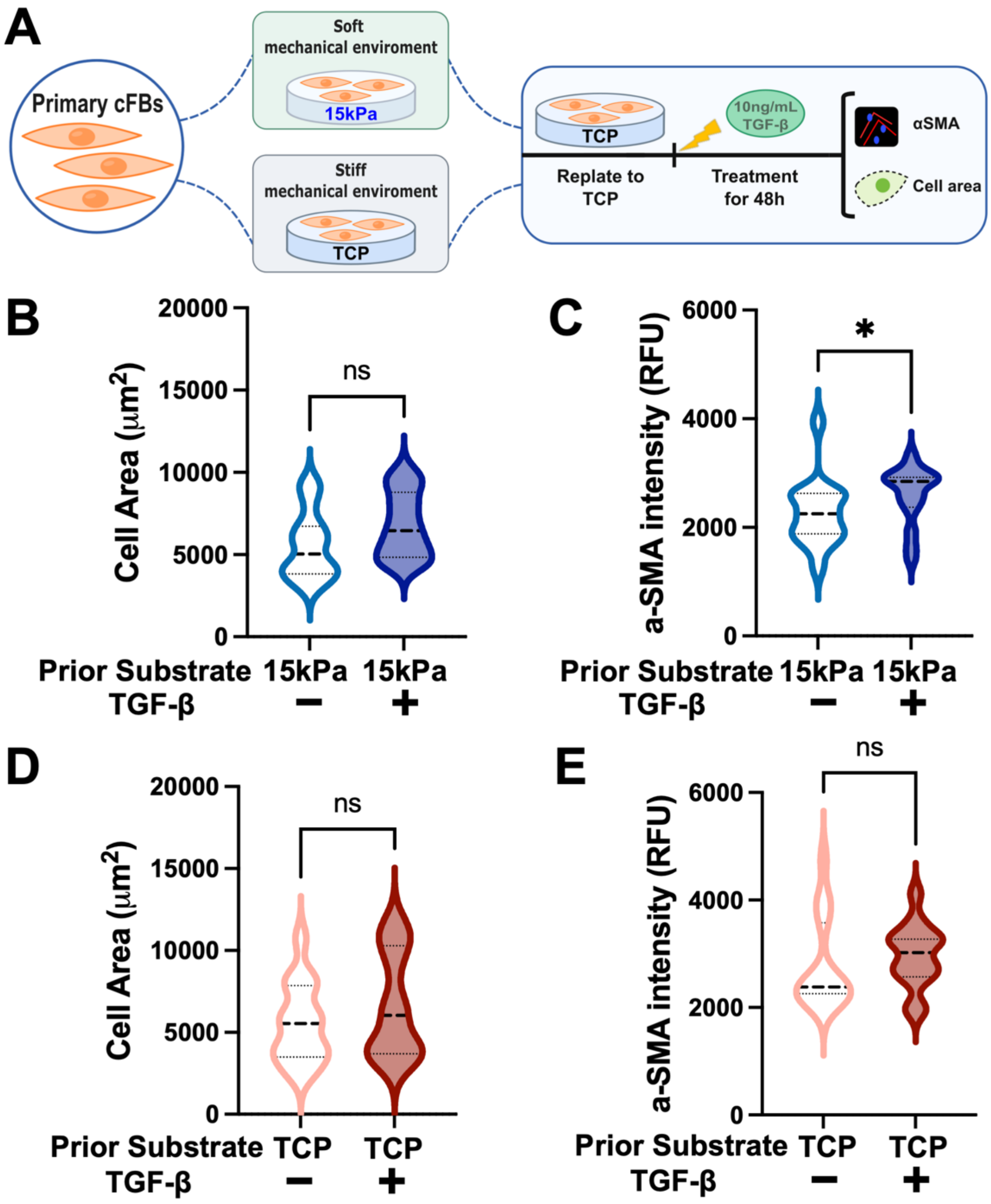


**Figure S2. Culture substrate affects TGF-β responsiveness of primary human cardiac fibroblasts.** **(A)** Schematic of experimental design showing primary cFBs maintained on 15kPa PDMS or TCP for two passages, replated onto TCP, and subsequently treated with TGF-β for 48 hr. **(B-C)** Quantification of **(B)** cell area and **(C)** α-SMA fluorescence intensity of primary cFBs cultured on 15kPa PDMS substrates left untreated or exposed to 10 ng/mL TGF-β. **(D-E)** Quantification of **(D)** cell area and **(E)** α-SMA fluorescence intensity of primary cFBs cultured on TCP left untreated or treated with 10 ng/mL TGF-β. Error bar: *SD*. ∗ indicate *p* value less than 0.05.

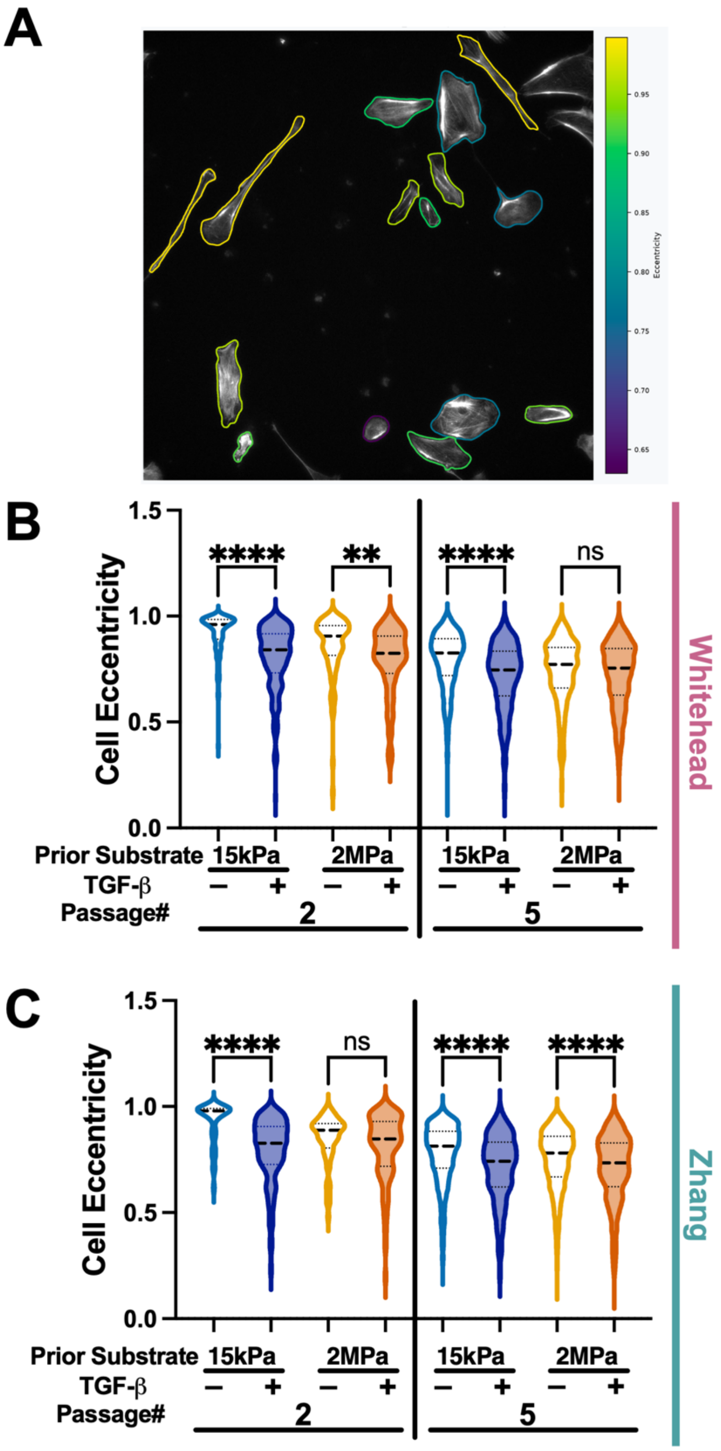


**Figure S3. Mechanical culture history and TGF-β modulate cell eccentricity in iPSC-derived cardiac fibroblasts. (A)** Representative image illustrating cell eccentricity analysis. Colored outlines delineate individual cells, with colors corresponding to eccentricity values ranging from 0 (circular) to 1 (highly elongated). **(B-C)** Quantification of cell eccentricity in **(B)** Whitehead- and **(C)** Zhang-derived MYBPC3fs iPSC-cFBs previously maintained on 15kPa or 2MPa PDMS substrates through P2 or P5, subsequently replated onto TCP and treated with or without 10 ng/mL TGF-β for 48 hr. Data were obtained from ≥ 3 independent experimental batches. Error bars represent SD. **** indicate $p$ values less than 0.0001.

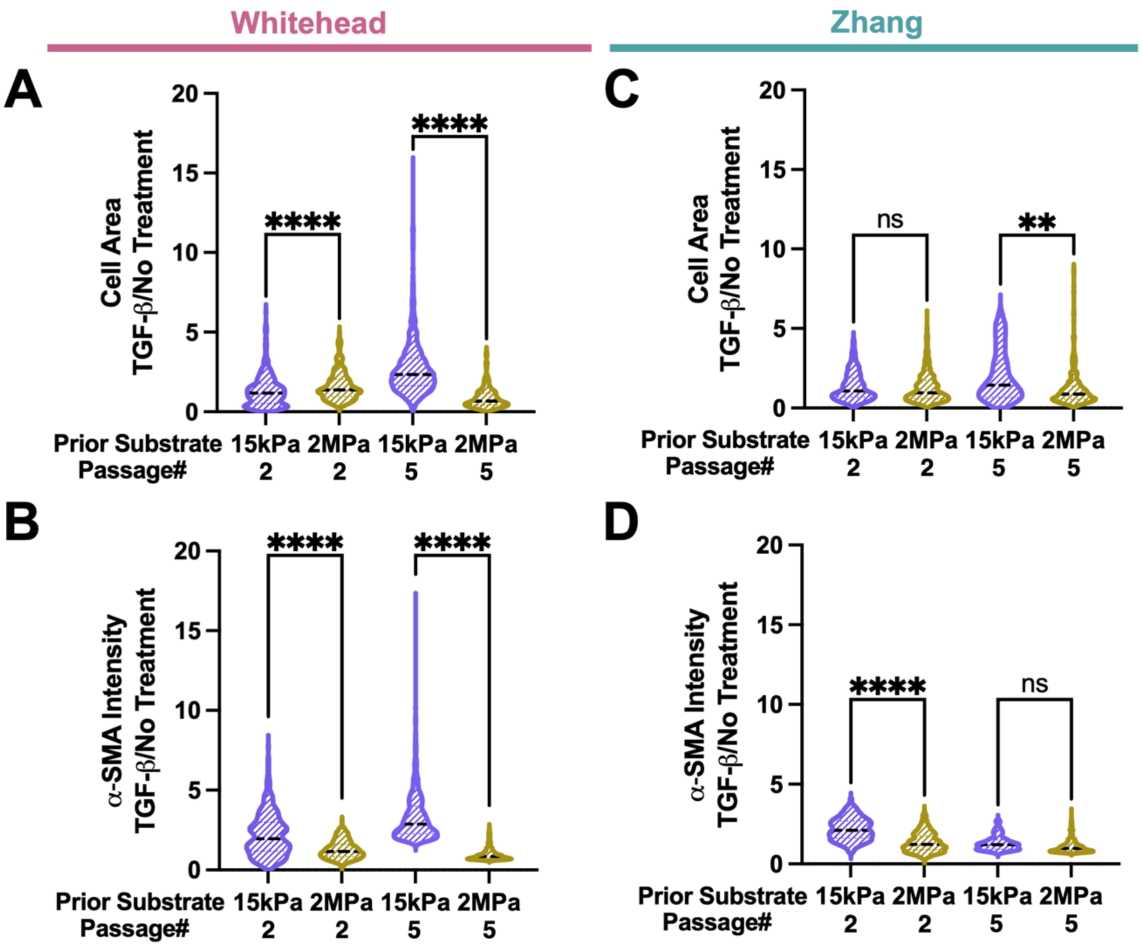


**Figure S4. Mechanical culture history and passage influence the relative TGF-β response of iPSC-derived cardiac fibroblasts in 2D culture**. **(A-B)** TGF-β treatment/no-treatment ratios of **(A)** cell area and **(B)** α-SMA fluorescence intensity in Whitehead-derived MYBPC3fs iPSC-cFBs previously maintained on 15kPa or 2MPa PDMS substrates through P2 or P5. **(C-D)** Corresponding TGF-β treatment/no-treatment ratios of **(C)** cell area and **(D)** α-SMA fluorescence intensity in Zhang-derived MYBPC3fs iPSC-cFBs across the same mechanical-history and passage conditions. Following mechanical conditioning, cells were replated onto TCP and treated with or without 10 ng/mL TGF-β for 48 hr. Ratios were calculated as the treatment-associated response relative to the corresponding untreated condition. n values represented ≥ 3 independent batches. Error bar: *SD*. $*$, $***$, $****$ indicate *p* value less than 0.05, 0.001 and 0.0001, respectively.

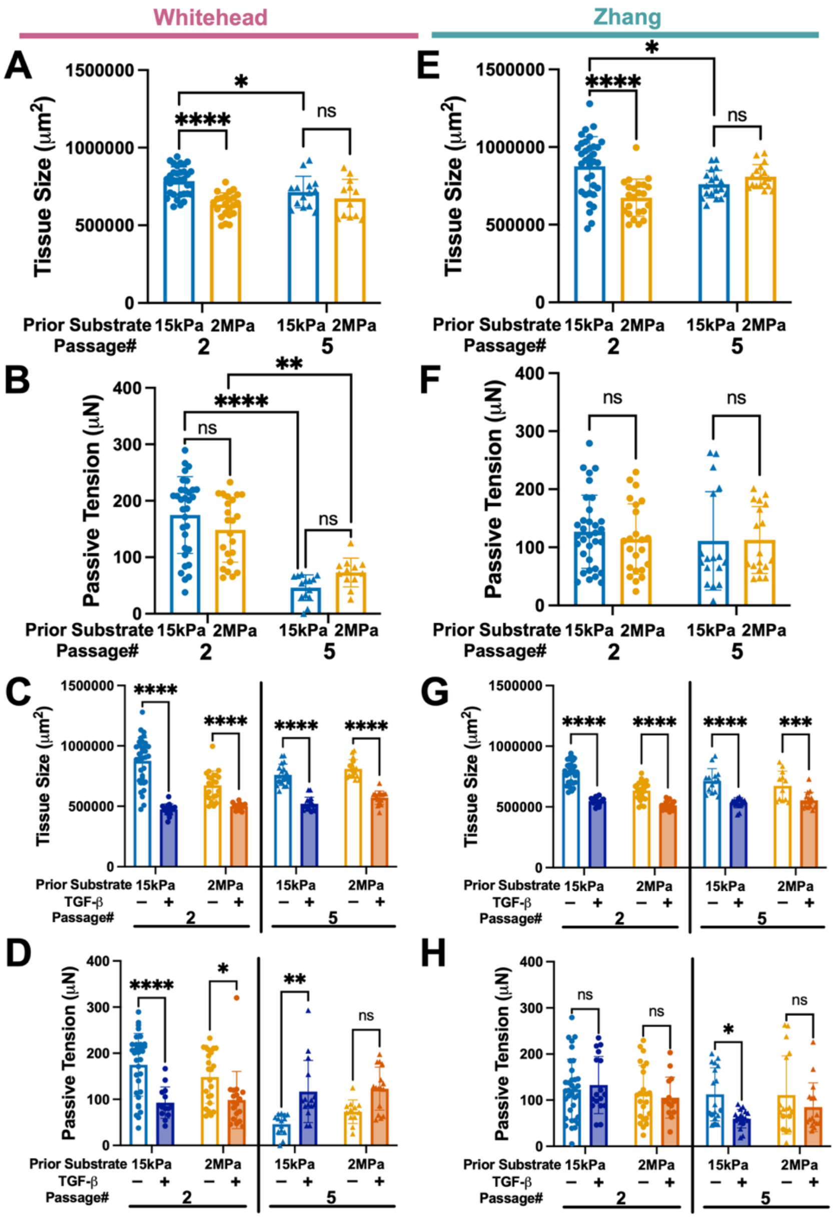


**Figure S5. Prior mechanical culture history and TGF-β influence the morphology and passive mechanics of fibroblast-only 3D tissues. (A-D)** Quantification of **(A)** basal tissue size, **(B)** basal passive tension, **(C)** tissue size following treatment with or without TGF-β, and **(D)** passive tension following treatment with or without TGF-β in fibroblast-only tissues generated from Whitehead-derived MYBPC3fs iPSC-cFBs previously maintained on 15kPa or 2MPa PDMS substrates through P2 or P5. **(E-H)** Corresponding quantification of **(E)** basal tissue size, **(F)** basal passive tension, **(G)** tissue size following treatment with or without TGF-β, and **(H)** passive tension following treatment with or without TGF-β in tissues generated from Zhang-derived MYBPC3fs iPSC-cFBs across the same mechanical-history and passage conditions. Tissues were treated with or without 10 ng/mL TGF-β1 from day 2 to day 10 following tissue formation. n values represented ≥ 3 independent batches. Error bar: *SD*. *, **, ***, **** indicate *p* value less than 0.05, 0.01, 0.001 and 0.0001, respectively.

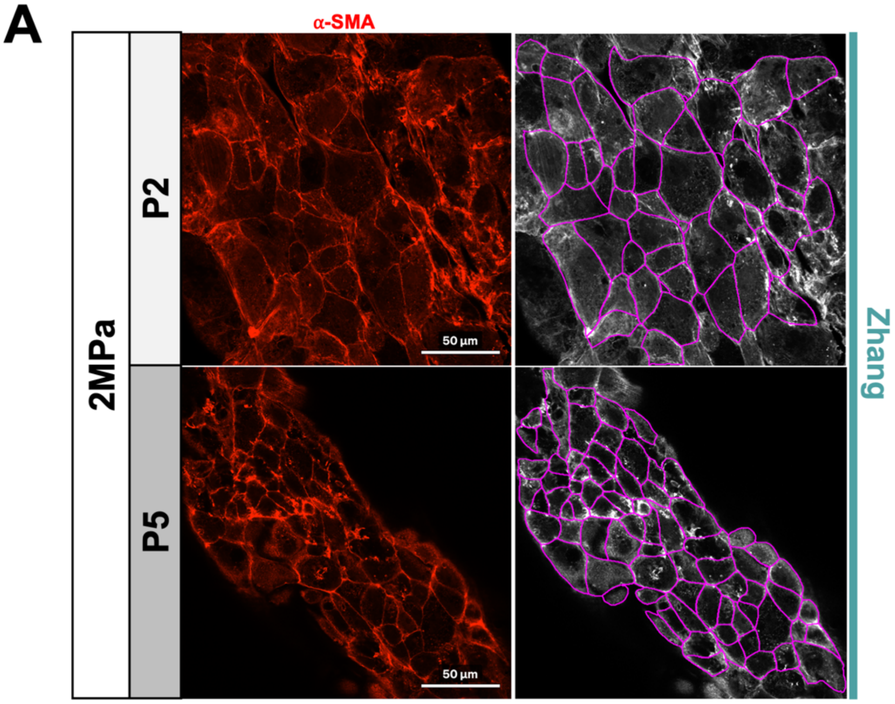


**Figure S6. Representative image-analysis workflow for quantification of cell area and α-SMA fluorescence intensity within fibroblast-only 3D tissues. (A)** Representative α-SMA fluorescence images (red, left) and corresponding cell-segmentation results (right) from tissues generated using Zhang-derived MYBPC3fs iPSC-cFBs previously maintained on 2MPa PDMS through P2 or P5. Magenta contours delineate individual cell boundaries identified by the image-analysis software and used for cell-area and fluorescence-intensity measurements. Scale bars, 50 µm.

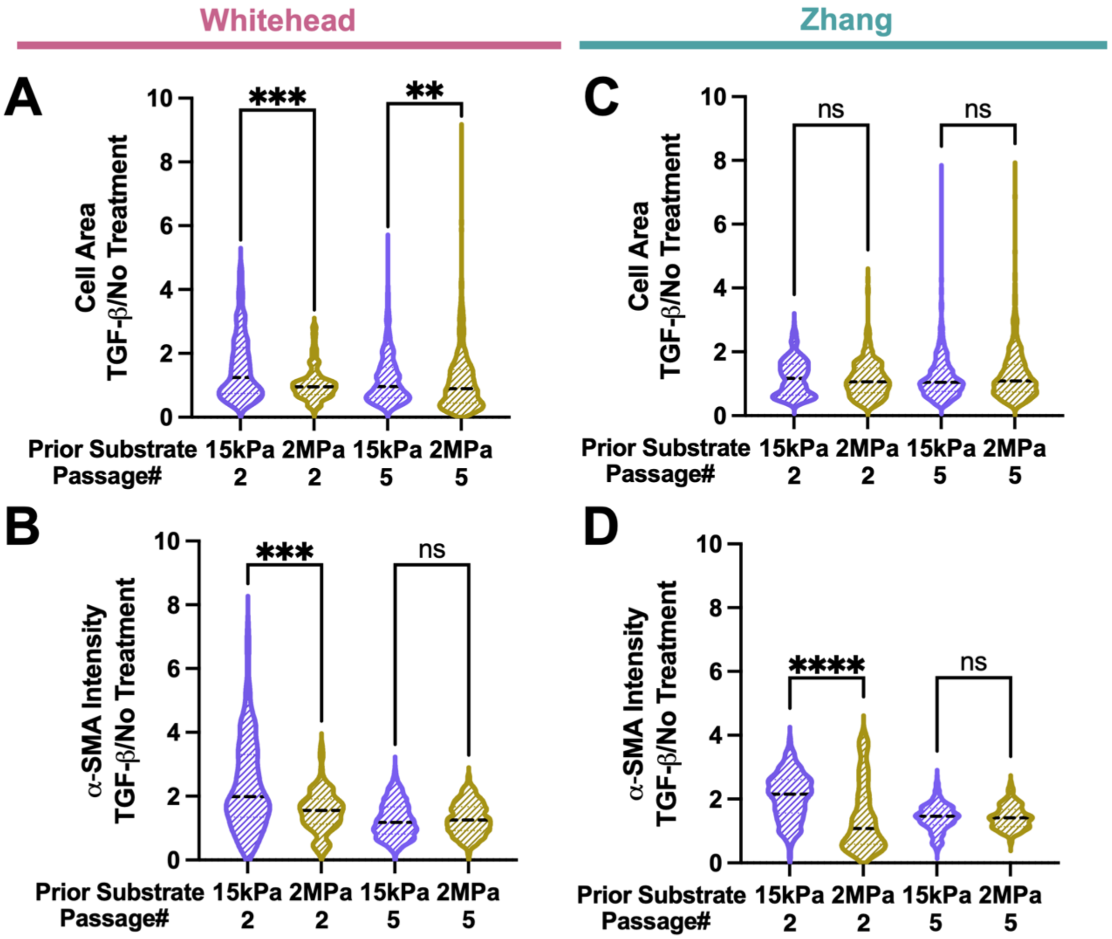


**Figure S7. Mechanical culture history and passage influence the relative cellular response to TGF-β within fibroblast-only 3D tissues. (A-B)** TGF-β treatment/no-treatment ratios of **(A)** cell area and **(B)** α-SMA fluorescence intensity within tissues generated from Whitehead-derived MYBPC3fs iPSC-cFBs previously maintained on 15kPa or 2MPa PDMS substrates through P2 or P5. **(C-D)** Corresponding TGF-β treatment/no-treatment ratios of **(C)** cell area and **(D)** α-SMA fluorescence intensity within tissues generated from Zhang-derived MYBPC3fs iPSC-cFBs across the same mechanical-history and passage conditions. Following tissue formation, constructs were cultured with or without 10 ng/mL TGF-β1 from day 2 to day 10. Ratios were calculated relative to the corresponding untreated condition. n values represented $\geq$ 3 independent batches. Error bar: *SD*. ∗, ∗∗, ∗∗∗, ∗∗∗∗ indicate *p* value less than 0.05, 0.01, 0.001 and 0.0001, respectively.
